# Generation and validation of an Acan-Cre mouse line to selectively label Class-B excitatory neurons of the cerebellar nuclei

**DOI:** 10.64898/2026.05.21.726923

**Authors:** Julian Cheron, Maggie Lowman, Manjari M-G Anant, Max Siauw, Justus M. Kebschull

## Abstract

The cerebellar nuclei form the main output structures of the cerebellum and are composed of a deeply conserved set of cell types. Two excitatory cell classes, Class-A and -B, are present in each cerebellar nucleus and mediate all excitatory output of the cerebellum. To provide genetic access to these cell types, here we identified *Acan* as a marker gene for Class-B cells and generated a knock-in Acan-P2A-Cre mouse line. We demonstrate that this Acan-Cre line selectively labels Class-B neurons in the cerebellar nuclei and validate its use in viral projection tracing. This new mouse line provides a valuable genetic tool to study cerebellar nuclei organization and function.

## Introduction

The cerebellar nuclei form the primary output structures of the cerebellum and play a central role in motor and cognitive functions (Kebschull et al., 2023; Rudolph et al., 2023; Uusisaari and De Schutter, 2011). A recent study revealed an unexpected diversity of neuronal subtypes within the cerebellar nuclei, including two classes of excitatory neurons---Class-A and Class-B---three classes of inhibitory neurons, and various non-neuronal cell types (**Fig. 1A**). Class-A and -B cells are responsible for all excitatory cerebellar output, shared across all cerebellar nuclei, and conserved across 300 million years of amniote evolution (Kebschull et al., 2020). These neurons thus occupy a critical circuit position as the cells integrating cerebellar computation and transmitting it to the rest of the brain. Each excitatory neuron class exhibits distinct projection patterns, morphology, and electrophysiological properties (Fujita et al., 2020; Kebschull et al., 2020; Uusisaari et al., 2007), and the two classes are hypothesized to differentially contribute to cerebellar function and disease as fundamentally distinct output channels of the cerebellum. In particular, Class-B cells map onto the large excitatory cells of the cerebellar nuclei and the previously reported GadNL cells of the mouse (Uusisaari et al., 2007), and are specifically expanded at the expense of Class-A cells in the human lateral nucleus (Kebschull et al., 2020). Large glutamatergic Class-B neurons may also be differentially vulnerable in neurodegenerative conditions such as Friedreich’s Ataxia, where large excitatory neurons of the lateral nucleus are preferentially affected (Koeppen et al., 2011). Despite their importance, however, no genetic tools exist to provide selective access to these cell types, hindering their functional dissection.

**Figure 1.**
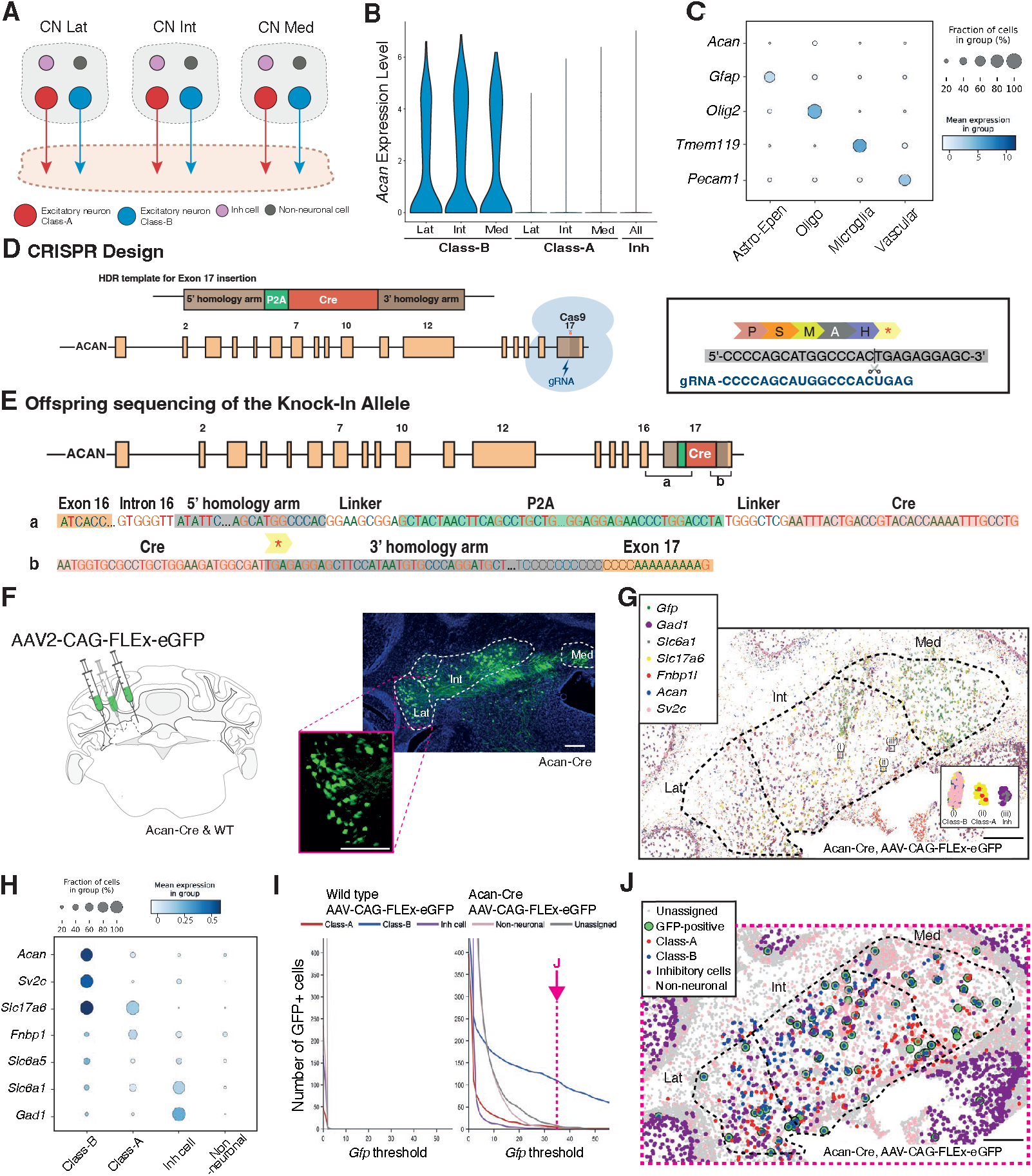
Generation and validation of the Acan-Cre mouse line. **(A)** Schematic overview of cell types, including Class-A, Class-B, inhibitory (Inh) neurons, and non-neuronal cells in the in the lateral (lat), interposed (int), and medial (med) cerebellar nuclei. **(B)** Expression of *Acan* mRNA across neuronal populations in the lat, int, and med cerebellar nuclei in the different neuronal cells. *Acan* expression is restricted to Class-B excitatory neurons. Data adapted from (Kebschull et al., 2020). **(C)** *Acan* expression in the cerebellum across non-neuronal cell populations from the Allen Brain Cell (ABC) Atlas, including astrocyte-ependymal (Astro-Epen), oligodendrocytes (Oligo), microglia, and vascular cells. *Acan* expression is minimal across all non-neuronal cell types. **(D)** Endogenous *Acan* locus and CRISPR/Cas9 targeting strategy. A P2A-Cre cassette was inserted at the 3’ end of the coding sequence to preserve endogenous *Acan* expression. Schematic of the HDR donor construct containing 5’ and 3’ homology arms flanking the P2A-Cre sequence. **(E)** Schematic representation of the knock-in allele showing the full *Acan* gene structure after insertion of the P2A-Cre sequence. (a, b) Validation of the targeted allele by long-read sequencing, confirming precise insertion at the *Acan* locus and sequence integrity. **(F)** Experimental design: injection of AAV2-CAG-FLEx-eGFP into the cerebellar nuclei of wild-type (WT) and Acan-Cre mice. Scale bar: 200 µm **(G)** In situ sequencing of multiple marker genes in the cerebellar nuclei, enabling classification of major neuronal and non-neuronal cell types. Insets (i), (ii), and (iii) show magnified views of a Class-A, a Class-B, and an inhibitory cell, respectively. Scale bar: 200 µm **(H)** Cell-type classification based on spatial transcriptomics data from (G), using combinatorial gene expression thresholds. **(I)** Quantification of *Gfp*-positive cells across a continuous range of *Gfp* thresholds for each identified cell-type category. *Gfp* expression remains strongly enriched in Class-B neurons across thresholds, with minimal labeling in other neuronal and non-neuronal populations. **(J)** Spatial overlay of cell-type classification and *Gfp*-positive cells (threshold = 35), showing selective labeling of Class-B excitatory neurons in Acan-Cre; AAV-CAG-FLEx-eGFP mice. Scale bar: 200 μm

Aggrecan, encoded by the *Acan* gene, is a major structural proteoglycan in cartilage, where it plays a critical role in skeletal development and resistance to mechanical load (Ahmed et al., 2024; Gibson and Briggs, 2016; Lin et al., 2021). In the central nervous system, Aggrecan is a major component of perineuronal nets, extracellular matrix structures that regulate neuronal plasticity (Rowlands et al., 2018). Manipulation of *Acan* expression can profoundly alter circuit dynamics, including reinstating juvenile-like plasticity in the adult visual cortex (Rowlands et al., 2018) and regulating synaptic plasticity in excitatory neurons of the hippocampal CA2 area (Noguchi et al., 2017). While conditional *Acan* loss-of-function models have been developed, no faithful Cre driver line is available to selectively target *Acan*-expressing neuronal populations (Rashid et al., 2017).

Here we describe an Acan-P2A-Cre knock-in mouse line on the C57Bl6/J background. We validate the specificity of this mouse line for Class-B neurons in the cerebellar nuclei and demonstrate that Class-B neurons exhibit distinct projection patterns compared to the aggregate of all excitatory neurons of the lateral cerebellar nucleus. This new mouse line will enable refined interrogation of cerebellar output circuits and offers broader applicability for studying other Acan-expressing populations and *Acan* gene function.

## Materials and Methods

### Animals

We conducted all procedures in accordance with the Johns Hopkins University Animal Care and Use Committee (ACUC) protocol MO23M346. We purchased C57Bl6/J and Slcl7a6-Cre (JAX stock #016963, (Vong et al., 2011)) mice from the Jackson Laboratory.

Acan-Cre knock-in mice were generated using CRISPR/Cas9-mediated genome editing targeting the endogenous *Acan* locus by the Johns Hopkins Transgenic Mouse Core Facility. Briefly, a P2A-Cre cassette was inserted immediately upstream of the endogenous stop codon of the Acan-201 transcript to preserve the *Acan* coding sequence while enabling co-expression of Cre recombinase (**Fig. 1D**). A guide RNA targeting the stop codon region was selected (CCCCAGCATGGCCCACTGAG), and a donor DNA construct containing the P2A-Cre cassette flanked by 5’ and 3’ homology arms was synthesized (GenScript; sequences are available in **Supplemental Data S1**). Genome editing was performed by pronuclear injection of CRISPR/Cas9 reagents and donor DNA into fertilized mouse oocytes (B6SJL background), followed by embryo transfer into pseudopregnant females (Wang et al., 2013; Yang et al., 2013). Founder animals were screened for correct insertion using PCR with primers that spanned the junction between genomic DNA and the inserted sequence on both sides of the targeted locus. An additional primer pair targeting Cre recombinase was used to confirm the presence of the transgene. Expected amplicon sizes are 100 bp for Cre, and 529 bp and 300 bp for the left and right targeted alleles, respectively. Positive founders were identified and bred to wild-type C57Bl6/J mice to establish germline transmission of the knock-in allele.

#### Genotyping Primer sequences

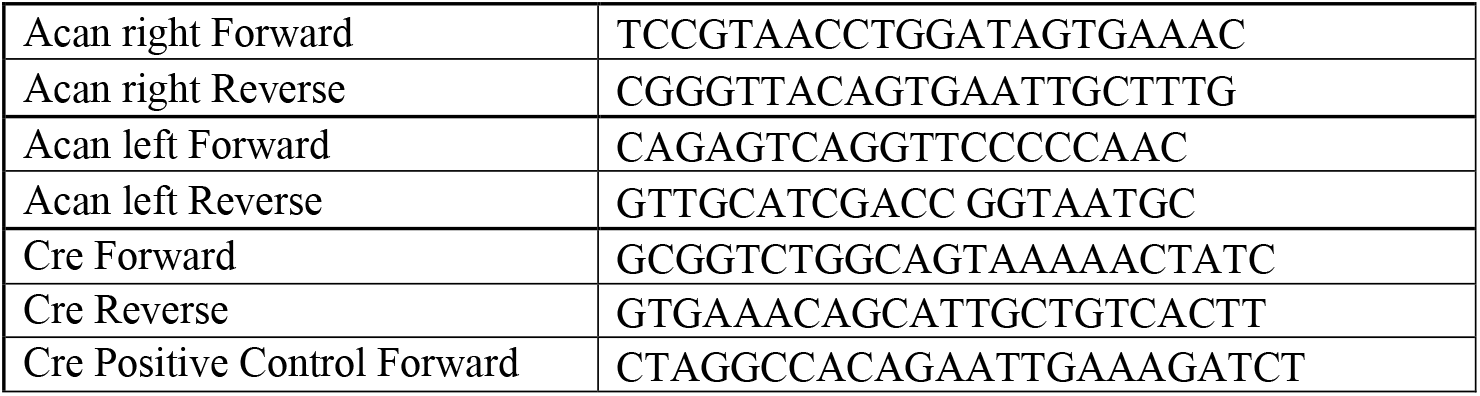
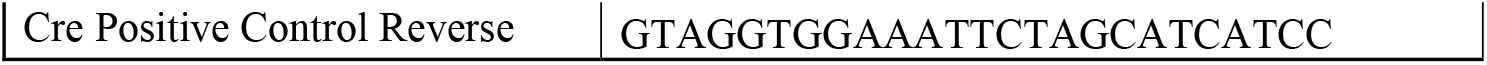

We used 8-12-week-old male mice for all experiments.

To confirm the correct insertion of the P2A-Cre cassette, we performed PCR amplification using the following primers and sequenced the PCR product by 0xford Nanopore (Plasmidsaurus).

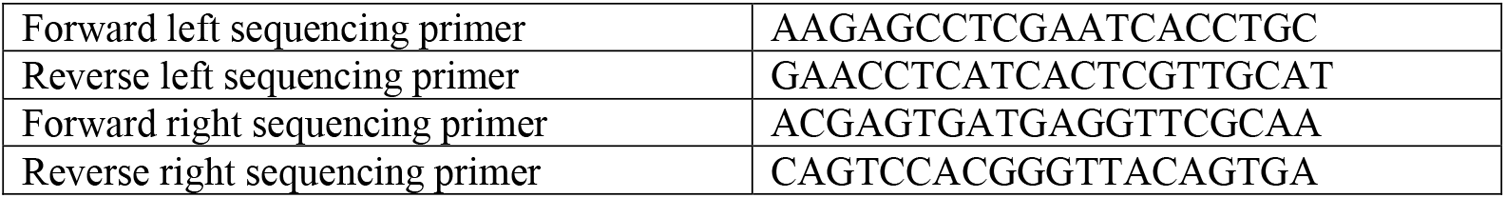

### Viral injections

We performed stereotaxic surgeries and viral injections using standard procedures. Briefly, we anesthetized mice using 1-2% isoflurane and placed them in a stereotaxic apparatus (Kopf Instruments). We pressure injected AAV virus into specific brain regions at a rate of 2 nL/second. For in situ transcriptomic profiling [N = 2 Acan-Cre mice and 1 wild-type (WT) mouse; **Fig. 1F-J**] and anterograde tracing (N = 5 S1c17a6-Cre mice and N = 7 Acan-Cre mice; **Fig. 2**) we injected 500 nL AAV2-CAG-FLEx-eGFP (Addgene #51502-AAV2 at 1.1×10^12^ vg/mL) at each site. We used the following coordinates in the right hemisphere (all relative to lambda); anterior medial nucleus: −1.83 mm AP, 0.75 mm ML, 3.4 mm DV; posterior medial nucleus: −2.18 mm AP, 1.0 mm ML, 3.2 mm DV; interposed nucleus: −1.83 mm AP, 1.5 mm ML, 3.3 DV; lateral nucleus: −1.4 mm AP, 2.45 mm ML, 3.8 mm DV. For in situ transcriptomics we targeted all 4 sites; for anterograde tracing we targeted the lateral cerebellar nucleus only. Animals were sacrificed 4 weeks after viral injection to allow sufficient viral expression.

**Figure 2.**
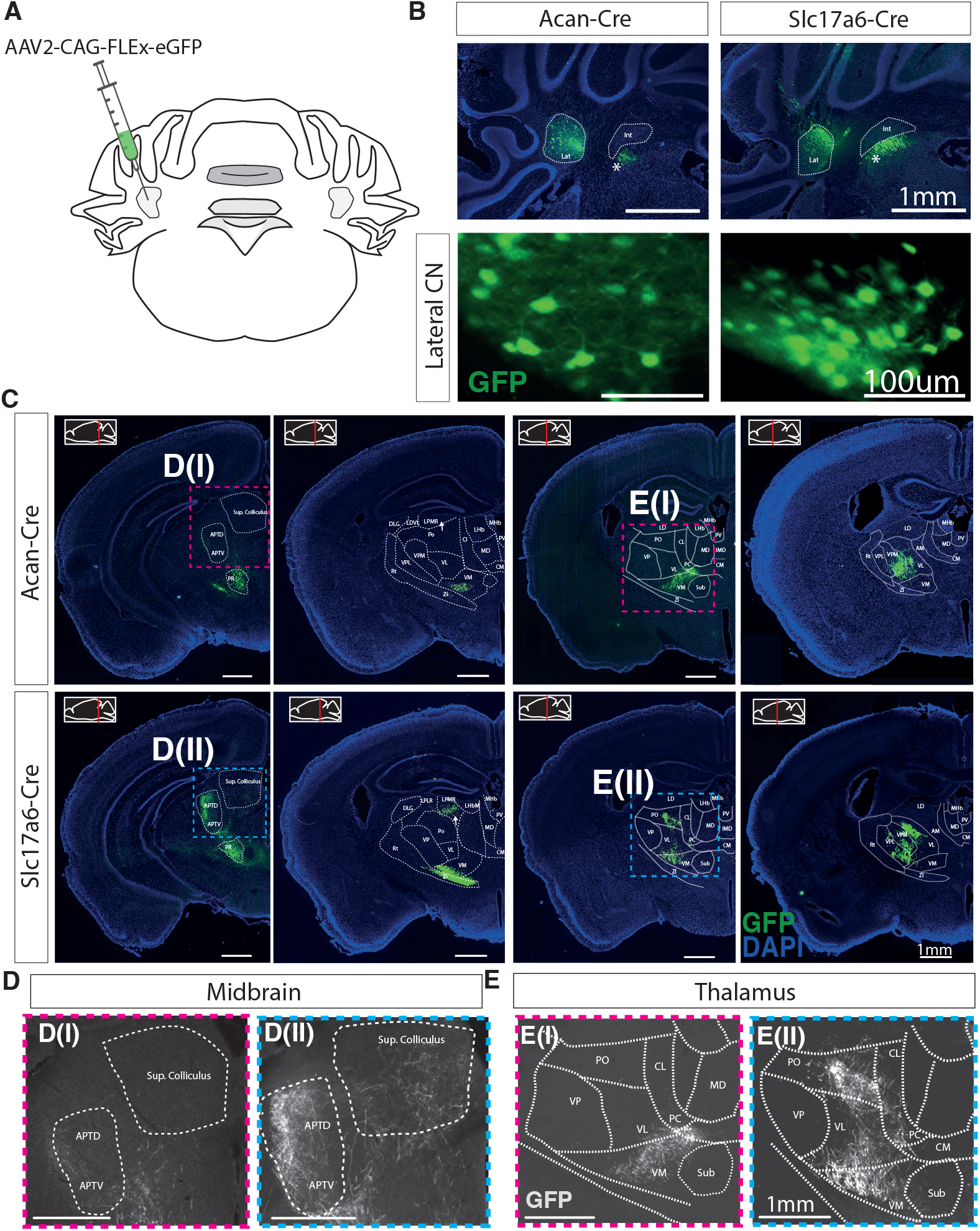
Projection patterns of Class-B neurons from the lateral cerebellar nucleus show selective connectivity. **(A)** Experimental design: injection of AAV2-CAG-FLEx-eGFP into the lateral cerebellar nucleus of Acan-Cre (n = 7) and S1c17a6-Cre (n = 5) mice. **(B)** Representative images show GFP-positive cells in the lateral cerebellar nucleus of Acan-Cre and S1c17a6-Cre mice. * indicates labeled axons. **(C)** Representative coronal sections illustrating axonal projections following AAV2-FLEx-eGFP injections in Acan-Cre and S1c17a6-Cre mice across selected target regions. Colored boxes indicate areas selected for higher-magnification views. First column: regions such as the superior colliculus (SC) and anterior pretectal nucleus (APT) are preferentially targeted by the complete excitatory population but show minimal or no innervation from Class-B neurons. Second column: the zona incerta (ZI) and ventromedial thalamic nucleus (VM) are targeted in both conditions, with ZI being more strongly targeted in S1c17a6-Cre mice than in Acan-Cre mice, whereas the lateral posterior thalamic region (LPMR) is selectively innervated in S1c17a6-Cre mice but not in Acan-Cre mice. Third column: the posterior (PO) thalamic nucleus is strongly innervated by S1c17a6-Cre neurons but not by Acan-Cre neurons. In contrast, nuclei including VM, VL, PC, and CM are similarly targeted in both lines. Fourth column: Acan-Cre projections form a spatially restricted subset within the broader projection fields of S1c17a6-Cre neurons, particularly within VP, VL, and VM.**(D-E)** Higher-magnification views of the boxed regions in C.

### BARseq3 in situ sequencing

In situ transcriptomic profiling was performed using BARseq3 in situ sequencing as previously described (Qi, Anant, Faltine-Gonzalez et al., 2026). Briefly, AAV-injected WT and Acan-Cre brains were extracted and fresh frozen. The cerebellar nuclei where sectioned at 16 pm thickness using a cryostat and mounted onto silanized and PDL coated glass-bottom dishes. Sections were fixed in 4% paraformaldehyde, permeabilized using cold methanol, and processed for BARseq3 library preparation as described. Gene-specific SNAIL probes were designed based on prior single-nucleus RNA sequencing data to enable discrimination of major neuronal populations (Kebschull et al., 2020). We targeted 11 marker genes (10 nM per probe in hybridization solution): *Acan, Fnbpll, Gfp, Gad1, Gfap, Slc17a6, Slc6al, Slc6a5, Sv2c, Pdgfra*, and *Tmem119*. We used 8 probe pairs per gene for *Acan, Sv2c, Slc6a5*, and *Fnbpll*, and 4 probe pairs per gene for the rest. In addition, we included SNAIL probes against *Gfp* to detect viral gene expression (0.1 nM per probe in hybridization solution; probe sequences are available in **Supplemental Data S2**). Following probe hybridization, samples underwent ligation and rolling circle amplification to generate spatially localized amplicons that were crosslinked to the tissue. Gene IDs were then determined by Illumina sequencing using a spinning-disk confocal microscope (Crest X-Light V3) mounted on a Nikon Ti2-E with a 40x water-immersion objective (NA 1.1). Image processing and gene calling was performed as previously described to generate a cell x gene matrix (Qi, Anant, Faltine-Gonzalez et al., 2026). Cell identity was then assigned by marker gene expression: Non-neuronal populations (including astrocytes, microglia, and oligodendrocytes) were identified based on established marker combinations (e.g., *Gfap, Tmem119, Pdgfra*) and the absence of neuronal signatures.

Excitatory neuronal subclasses *(S1c17a6* positive cells) were resolved into Class-A and Class-B populations using differential expression patterns of *Fnbpll, Acan*, and *Sv2c*, while inhibitory neurons were identified based on *Gadl* and *Slc6al* expression profiles. Thresholds were empirically determined from gene expression distributions to ensure consistency with previously defined snRNA-seq cell type annotations. Cells expressing general neuronal markers but not satisfying any predefined combinatorial rule were conservatively assigned to an “unassigned” category.

### Anterograde tracing

For anterograde tracing (**Fig. 2**), we transcardially perfused AAV-injected mice with 1×PBS followed by 4% paraformaldehyde (PFA; Boster Biological Technology, AR1068). We dissected the brain, post-fixed it in 4% PFA at 4 °C overnight, and cryoprotected it in 30% sucrose (Millipore Sigma, S5016). We then embedded the brain in 0CT, froze it using methylbutane at −65 °C, and then stored the samples at −80 °C until cryosectioning. We cryosectioned the samples into 30 μm slices and mounted them onto Superfrost Plus Microscope Slides (Fisher Scientific, 1255015). Sections were immunostained using a chicken anti-GFP primary antibody (Invitrogen, A10262, 1:200) followed by a goat anti-chicken Alexa Fluor 488 secondary antibody (Invitrogen, A11039, 1:1000) and imaged on a BZ-X700 epifluorescence microscope (Keyence).

### Data and code availability

Analysis code used for cell-type classification and downstream analyses is available at: https://github.com/kebschulllab/Acan-Cre-cerebellar-nuclei

BARseq3 datasets generated in this study have been deposited in Zenodo under the following D0I: 10.5281/zenodo.20214220.

Acan-Cre mice are available upon request.

## Results

### Generation of an Acan-P2A-Cre knock-in mouse line

To select a marker gene for Class-A or -B cells that could be used to construct a Cre-driver mouse line, we queried published single-nucleus RNA sequencing data of NeuN+ neurons in the mouse cerebellar nuclei (Kebschull et al., 2020) as well as of non-neuronal cerebellar cells in the Allen Brain Cell (ABC) Atlas (Yao et al., 2023). Class-A and -B cells in the medial, interposed, and lateral cerebellar nuclei differentially express several genes, some specific to each nucleus, others specific to each class across all nuclei. Among these genes, we selected *Acan* as a promising marker for Class-B neurons. It is expressed at moderate levels in Class-B cells across all cerebellar nuclei, with no detectable expression in Class-A or inhibitory neurons and minimal expression in non-neuronal cells (**Fig. 1B,C**). *Acan* is thus a promising gene to drive recombinase expression specifically in Class-B cells. An Acan-creERT2 mouse line already exists (Jax stock #019148) (Henry et al., 2009) but unfortunately does not label cells in the cerebellar nuclei (data not shown), likely because of an unintended deletion in the *Acan* gene 3’UTR in this specific line (Rashid et al., 2017). We therefore set out to develop a new Acan-Cre driver line.

To drive Cre expression mimicking *Acan* expression levels without disrupting *Acan* expression, we chose to replace the endogenous *Acan* stop codon with a P2A-Cre cassette. To do so, we designed a CRISPR/Cas9 guide RNA targeting a sequence close to the endogenous stop codon of the *Acan* gene and an HDR donor including the P2A-Cre cassette flanked by 5’ and 3’ homology arms corresponding to the targeted locus (**Fig. 1D, Supplemental Data S1**). After injection into B6SJL oocytes and embryo transfer, we identified founder animals that successfully transmitted the modified allele to F1 progeny and backcrossed these animals onto a C57Bl6/J background for at least 5 generations. We confirmed correct integration of the knock-in cassette into the endogenous locus by PCR genotyping and long-read sequencing (**Fig. 1E**). Hemizygous animals are overtly healthy but homozygous mice exhibit a dwarf phenotype, indicating a partial disruption of *Acan* function. We therefore maintained the Acan-Cre line as hemizygous.

### Acan-Cre selectively labels Class-B excitatory neurons in the adult cerebellar nuclei

To determine whether our new Acan-Cre line provides selective access to Class-B neurons, we performed stereotaxic injections of a Cre-dependent AAV (AAV2-CAG-FLEx-eGFP) into the cerebellar nuclei of hemizygous Acan-Cre adult mice and wild type control mice (**Fig. 1F**). As expected, we observed strong viral labeling of large neurons in all three cerebellar nuclei only in Acan-Cre animals (**Fig. 1F**). We then used BARseq3 spatial transcriptomics to determine the molecular identities of the Gµ-labeled cells (Qi, Anant, Faltine-Gonzalez et al., 2026). In both wild type and Acan-Cre animals, we probed for *Gfp* mRNA and a panel of marker genes selected to distinguish major neuronal and non-neuronal populations of the cerebellar nuclei: excitatory neurons were identified based on expression of *S1c17a6 (Vglut2)*, a pan-glutamatergic marker labeling both Class-A and Class-B populations. Within this population, *Acan* and *Sv2c* expression were used to identify Class-B neurons, while *Fnbp1l* served as a marker enriched in Class-A neurons. Inhibitory neurons were identified using *Gadl, Slc6al* (GABAergic marker), and *Slc6a5* (glycinergic marker). The special population of rhombic-lip derived, large glycinergic cells (Bagnall et al., 2009) found in the medial nucleus was counted as Class-B neurons in accordance with Kebschull et al., 2020. Non-neuronal populations were identified using *Gfap* as an astrocytic marker, *Tmem119* as a microglial marker and *Pdgfra* as an oligodendrocyte marker. This marker combination enabled robust classification of cerebellar nuclei cell types (**Fig. 1G,H**) and allowed us to determine which cell types were labeled in the viral injections.

As expected in the dense neuropil of the brain and from high viral expression levels, *Gfp* signal was detected not only within neuronal somata but also in axons and dendrites in virus injected samples, which may contaminate gene calling in spatial data. Therefore, we evaluated the robustness of viral labeling of specific cell types across different *Gfp* expression thresholds (**Fig. 1I**). Across all thresholds, *Gfp* detection remained strongly enriched in Class-B neurons, whereas Class-A, inhibitory, and non-neuronal populations showed minimal labeling. Unassigned cells, which have low quality gene expression data, also displayed low levels of *Gfp* signal and likely reflect low quality cells (**Fig. 1I,J**). Taken together, these results demonstrate that Acan-Cre provides selective genetic access to the Class-B population in the adult cerebellar nuclei.

### Class-B excitatory neurons exhibit distinct projection patterns

With the Acan-Cre mouse line in hand, we then sought to contrast the brain-wide projection patterns of Class-A and -B neurons of the lateral nucleus. To this end, we injected Cre-dependent AAV (AAV2-CAG-FLEx-eGFP) into the lateral nucleus of either Acan-Cre or S1c17a6-Cre mice (**Fig. 2A**). As demonstrated above, viral injections into Acan-Cre animals label Class-B cells, whereas injections into S1c17a6-Cre mice label all excitatory cells and thus both Class-A and -B neurons. Projections found in S1c17a6-Cre animals, therefore, represent the total excitatory output of the lateral nucleus, and projections found in Acan-Cre animals should be a subset of these projections. Conversely, any brain regions innervated only in S1c17a6-Cre animals are likely exclusively innervated by A-class cells. Indeed, we found that projections in Acan-Cre animals are a subset of regions innervated in S1c17a6-Cre animals (**Fig. 2C-E**). Several regions, including the superior colliculus and the anterior pretectal nucleus (APT), were robustly innervated in S1c17a6-Cre mice but showed minimal or no innervation in Acan-Cre mice (**Fig. 2C,D**). Similarly, projections to the posterior and lateral posterior thalamic nuclei (PO and LPMR) were prominent in S1c17a6-Cre mice but largely absent in Acan-Cre mice (**Fig. 2C,E**). In contrast, thalamic nuclei such as VM, VL, PC, and CM were comparably targeted in both lines, indicating partial overlap in projection targets. ZI was targeted by both S1c17a6 and Acan projections but was more prominent for the S1c17a6 projection. Interestingly, we often found that in brain regions labeled in both mouse lines, including VP, VL, and VM, Acan-Cre projections formed spatially restricted subdomains embedded within the broader projection fields observed in S1c17a6-Cre mice (**Fig. 2C**, leftmost column), suggesting functional segregation within these target regions.

Together, these results demonstrate that Class-B neurons of the lateral nucleus exhibit selective and spatially restricted projection patterns within the broader excitatory cerebellar output system and validate the usefulness of our new Acan-P2A-cre line in understanding cerebellar output.

## Discussion

Here we introduce an Acan-P2A-Cre mouse line and validate it as a genetic strategy to give selective access to Class-B neurons in the cerebellar nuclei. This mouse driver line provides a critical entry point for studying this previously inaccessible excitatory neuron population in isolation. More broadly, our work provides a tool for interrogating Acan-expressing neuronal populations and the *Acan* function across the brain.

Initial work on the cellular composition of the cerebellar nuclei generally focused on morphology alone, distinguishing large, presumably excitatory, and small, presumably inhibitory, neurons. Since then, significant diversity has been discovered: Electrophysiological studies distinguished GABAergic and different non-GABAergic populations based on their morphological and electrophysiological properties (Uusisaari et al., 2007; Uusisaari and Knöpfel, 2011, 2008). Single-nucleus RNA sequencing then revealed molecular heterogeneity in excitatory cerebellar nuclei neurons, which mapped onto the previous electrophysiological distinctions (Kebschull et al., 2020). Based on these findings, we proposed a fundamental division of excitatory output from each cerebellar nucleus into Class-A and -B cells (Kebschull et al., 2023). Supporting this distinction, it was found that extensive projectional heterogeneity in excitatory cell types exists in the medial nucleus (Fujita et al., 2020), and indirect retrograde labeling approaches suggest distinct projection targets for Class-A and Class-B neurons of the lateral nucleus (Kebschull et al., 2020). Here, we now provide an independent and convergent assessment of this projection organization, further supporting the existence of distinct excitatory output circuitry within the cerebellar nuclei.

While the Acan-Cre line gives direct access only to Class-B cells, we expect that Class-A populations will be accessible with intersectional approaches; for example, in an S1c17a6-FlpO (Jax #0302l2) x Acan-Cre mouse cross, Class-A cells could be accessed with a FlpON; CreOFF construct (Fenno et al., 2017, 2014; Hughes et al., 2024). Such approaches may ultimately allow within-animal comparisons of the two major excitatory output channels of the cerebellar nuclei and facilitate causal dissection of their respective contributions to brain-wide network function.

## ACKNOWLEDGMENTS

This work was supported by the Klingenstein Simons and Sloan Fellowships to J.M.K., a Friedreich’s Ataxia Research Alliance Postdoctoral Fellowship to J.C., and a Kavli Distinguished PhD Fellowship to M.M.G.A..

## AUTHOR CONTRIBUTIONS

J.C. and J.M.K. conceptualized the study. J.C. and M.L. performed most of the mouse work. M.M.G.A. performed the BARseq3 experiment. J.C. performed all other experiments with help from M.S.. J.C. and J.M.K. wrote the manuscript with input from all authors.

## DECLARATION OF INTERESTS

We declare no competing interests.

## Abbreviations

SC: superior colliculus
APT: anterior pretectal nucleus
LP: lateral posterior thalamic nucleus
PO: posterior thalamic nucleus
VM: ventromedial thalamic nucleus
VL: ventrolateral thalamic nucleus
PC: paracentral thalamic nucleus
CM: centromedian thalamic nucleus
VP: ventral posterior thalamic nucleus
PR: prerubral field
MD: mediodorsal thalamic nucleus
CL: central lateral thalamic nucleus
MHb: medial habenula
LHb: lateral habenula
Rt: reticular thalamic nucleus
VPM: ventral posteromedial thalamic nucleus
VPL: ventral posterolateral thalamic nucleus
LD: laterodorsal thalamic nucleus
DLG: dorsal lateral geniculate nucleus
Sub: submedius thalamic nucleus
IMD: intermediodorsal thalamic nucleus
PV: paraventricular thalamic nucleus
ZI: zona incerta.

## Notes

### Competing Interest Statement

The authors have declared no competing interest.

