## Supplemental Data S1 for "Generation and validation of an Acan-Cre mouse line to selectively label Class-B excitatory neurons of the cerebellar nuclei"

**5’ homology arm:** ATATTCCGGAGCCCACAATGCACAGGGAGGGGCTTCTGTCCATGTGGGAAGGCTGCTCAGAGACTAAGCTTGGCCCTGCCTCCTCCCCTCTCACTGTTGATTTCCCACCTCCCTTTCTCGCTGCAGCCAACACCTACAAGCACAGGCTACAGAAGCGGAGCATGAGACCCACACGGAGGAGCCGCCCCAGCATGGCCCAC

**Linker-P2A-Linker-Cre:**

GGAAGCGGAGCTACTAACTTCAGCCTGCTGAAGCAGGCTGGAGACGTGGAGGAGAACCCTGGACCTATGGGCTCGAATTTACTGACCGTACACCAAAATTTGCCTGCATTACCGGTCGATGCAACGAGTGATGAGGTTCGCAAGAACCTGATGGACATGTTCAGGGATCGCCAGGCGTTTTCTGAGCATACCTGGAAAATGCTTCTGTCCGTTTGCCGGTCGTGGGCGGCATGGTGCAAGTTGAATAACCGGAAATGGTTTCCCGCAGAACCTGAAGATGTTCGCGATTATCTTCTATATCTTCAGGCGCGCGGTCTGGCAGTAAAAACTATCCAGCAACATTTGGGCCAGCTAAACATGCTTCATCGTCGGTCCGGGCTGCCACGACCAAGTGACAGCAATGCTGTTTCACTGGTTATGCGGCGGATCCGAAAAGAAAACGTTGATGCCGGTGAACGTGCAAAACAGGCTCTAGCGTTCGAACGCACTGATTTCGACCAGGTTCGTTCACTCATGGAAAATAGCGATCGCTGCCAGGATATACGTAATCTGGCATTTCTGGGGATTGCTTATAACACCCTGTTACGTATAGCCGAAATTGCCAGGATCAGGGTTAAAGATATCTCACGTACTGACGGTGGGAGAATGTTAATCCATATTGGCAGAACGAAAACGCTGGTTAGCACCGCAGGTGTAGAGAAGGCACTTAGCCTGGGGGTAACTAAACTGGTCGAGCGATGGATTTCCGTCTCTGGTGTAGCTGATGATCCGAATAACTACCTGTTTTGCCGGGTCAGAAAAAATGGTGTTGCCGCGCCATCTGCCACCAGCCAGCTATCAACTCGCGCCCTGGAAGGGATTTTTGAAGCAACTCATCGATTGATTTACGGCGCTAAGGATGACTCTGGTCAGAGATACCTGGCCTGGTCTGGACACAGTGCCCGTGTCGGAGCCGCGCGAGATATGGCCCGCGCTGGAGTTTCAATACCGGAGATCATGCAAGCTGGTGGCTGGACCAATGTAAATATTGTCATGAACTATATCCGTAACCTGGATAGTGAAACAGGGGCAATGGTGCGCCTGCTGGAAGATGGCGAT

**3’ homology arm:**

TGAGAGGAGCTTCCATAATGTGCCCAGGATGCTGAGCCCAGCGGCCAGCCAGGCTGACCGTGCATCCCACCCACATGGTGTCTTCTTGTCGCTTTTTGTCATATAAGGAATCCATTAAAGAAGGAAAAAAAACCCACATTGTGTGTGCCCCACTCCCCGCATCACCCAAACTGCATCTAATTTATCCCCTTCTCCCCCCC
